# Bone collagen hydrogen isotopes as tracers of local palaeoclimate: The Aurignacian from Isturitz cave (France) as a case study

**DOI:** 10.1101/2025.10.15.682524

**Authors:** M. Ibáñez-Herranz, L. Agudo-Pérez, M. Fernández-García, A. Cicero-Cabañas, M.C. Soulier, C. Normand, A.B. Marín-Arroyo

## Abstract

Hydrogen stable isotopes (δ^2^H) preserved in bone collagen have been proposed as a promising proxy for palaeoclimatic reconstruction, though empirical applications remain limited. To evaluate the reliability and scope of this approach, we performed a multi-isotopic analysis (δ^13^C, δ^15^N, δ^34^S, and δ^2^H) on 50 herbivore bone samples collected from three archaeostratigraphic layers spanning the Aurignacian sequence of Isturitz Cave (France). This archaeological well-preserved, dated and studied site provides a robust framework for testing isotopic proxies across temporal and taxonomic gradients. Our results reveal a progressive δ^2^H depletion over ∼3000-year interval, consistent with predicted increasing aridity. We also identified species-specific δ^2^H differences, underscoring the need for taxonomic resolution in future applications and in general palaeoclimatic interpretations. While these samples reflect faunal remains accumulated through human activity—introducing potential ecological and sampling biases—they nonetheless provide valuable records of past local conditions. Overall, our findings support the potential of bone collagen δ^2^H as a sensitive indicator of past hydrological conditions and a complementary tool for reconstructing both climatic trends and human-environment dynamics.

## 1. INTRODUCTION

Over the past few decades, geochemical techniques such as stable-isotope analyses have transformed the study of past ecosystems, establishing themselves as a valuable tool for determining palaeoecological and palaeoenvironmental conditions (Stevens et al., 2025 and references therein). Among the array of available possibilities, the analysis of δ^18^O and δ^13^C in dental enamel emerged as the prevailing approach for studying past climates. This method is of particular interest due to its capacity to investigate seasonality in herbivores through transects of phosphate or carbonate of dental hydroxyapatite. However, it is worth noting that these analyses provide insights on a time scale corresponding to the mineralisation of the tooth, which occurs during the first years of the animal’s life (Balasse, 2002; Hoppe et al., 2004; Kohn, 2004). In an attempt to address this deficit, some studies have examined the extraction of this isotope from bone bioapatite, though it appears not to be as reliable on bone tissues due to diagenetic alterations (Lee-Thorp, 2008; Pestle et al., 2014). Consequently, the door is open to exploring alternative approaches to studying past climates directly on this kind of biological remains recovered from archaeological and palaeontological sites. Among all the analysable biological materials, bone collagen is attracting considerable attention for its relative abundance and preservation at archaeological sites and its capacity to provide insights into behavioural, dietary, and even climatic information over a presumably longterm period during the lifetime of an animal (Matsubayashi and Tayasu, 2019; Quinn, 2024).

The isotopes most often studied in this context and material type are δ^13^C and δ^15^N, which offer valuable information about an animal’s diet and habitat, and in turn its palaeoenvironmental setting. Carbon values are highly correlated with plant metabolism, which varies depending on the habitat’s resources and environmental conditions, differing between C3 and C4 photosynthetic pathways (Van der Merwe, 1982; Van Klinken et al., 1994). Thus, based on the principle of isotopic accumulation through the food chain, animal collagen values reflect, to a certain extent, the ecological characteristics of the plants they consumed, where lower values would indicate colder and wetter conditions (Ehleringer and Monson, 1993; Kohn, 2010; Tieszen, 1991). Similarly, an analogous correlation has been observed between the nitrogen isotopic values of bone collagen and those of plants (Hartman, 2011; Reade et al., 2023). The latter depend on soil conditions and biological factors, such as mycorrhizal symbiosis, and are subsequently influenced by additional variables, including temperature and humidity (Craine et al., 2009). This results in more arid systems being categorised by higher values of δ^15^N (Craine et al., 2015; Heaton et al., 1986). Furthermore, another stable isotope that has attracted attention in this field is δ^34^S, which has also been utilised to deduce dietary and ecological information. In particular, sulphur appears to be especially useful for evaluating habitat use regarding mobility and migratory behaviours, given its strong reflection of geological influences and marine spray (Drucker et al., 2011; González-Rabanal et al., 2025; Jones et al., 2019; Privat et al., 2007).

However, two stable isotopes remain underutilised and that could be a direct reflection of climatic conditions, ^1^H and ^2^H. Also known as protium and deuterium, the relation between them (δ^2^H) has been considered a potential proxy for studying climates on organic tissues for decades (Cormie et al., 1994; Epstein et al., 1977; Estep and Dabrowski, 1980). The primary sources of hydrogen in animals’ tissues of terrestrial ecosystems are the meteoric water they drink or that is incorporated into lower levels of the food chain, making it a more straightforward source for studying precipitation than with other isotopes (Cormie et al., 1994; Hobson et al., 1999; Topalov et al., 2019; Tuross et al., 2008). Meteoric deuterium values are of particular interest due to their global variation, which is attributable to the strong isotopic fractionation resulting from condensation and precipitation processes inherent to the water cycle. Moreover, precipitation values depend on a series of geospatial characteristics, including altitude, latitude and distance from the sea; as well as on climatological or meteorological factors such as aridity, relative humidity or temperature; this results in a locally specific variation of δ^2^H (Bowen et al., 2005; Bowen and West, 2019; Kendall and Doctor, 2003; Kern et al., 2020; Pfahl and Sodemann, 2014; Vystavna et al., 2021).

Indeed, several fields have exploited δ^2^H on organic tissues to infer migration patterns, trophic levels, seasonal changes and even climatic conditions (Bearhop et al., 2003; Cryan et al., 2004; Dufour et al., 2025; Eerkens et al., 2020; Estep and Dabrowski, 1980; Hobson et al., 2012; Ruokonen et al., 2019; Sharp et al., 2003; White et al., 1994). Despite all of this potential, the main reason why deuterium has been overlooked in bone collagen research is due to the difficulty of obtaining direct and reliable values of isotopic carbon-bonded hydrogens, because a fraction of them is easily exchangeable with environmental water or vapour (DeNiro and Epstein, 1981). However, this problem has already been overcome analytically (Sauer et al., 2009; Soto et al., 2017; Wassenaar et al., 2023). Of the few studies that have analysed archaeological bone collagen, many have focused on palaeodietary or palaeoecology reconstructions (e.g., Arnay-de-la-Rosa et al., 2010; Gröcke et al., 2017; Reynard and Hedges, 2008; Topalov et al., 2013). From a climatological perspective, Leyden et al. (2006) and Reynard et al. (2020) found consistency of δ^2^H from bone collagen being related to precipitation, which confirms its potential as a palaeoclimatic proxy (*Figure 1*). Nevertheless, they were based on largescale geographic regions such as North America or the Mediterranean region, where multiple factors can influence the interspecific and interpopulation values of the samples. Certainly, some studies have noted that non-exchangeable hydrogen values measured in mammalian tissues appear to vary between species (Reynard et al., 2020; Reynard and Hedges, 2008; Topalov et al., 2013), hypothesising that this is due to the different body sizes and their inherent different water intake requirements.

**Figure 1.**
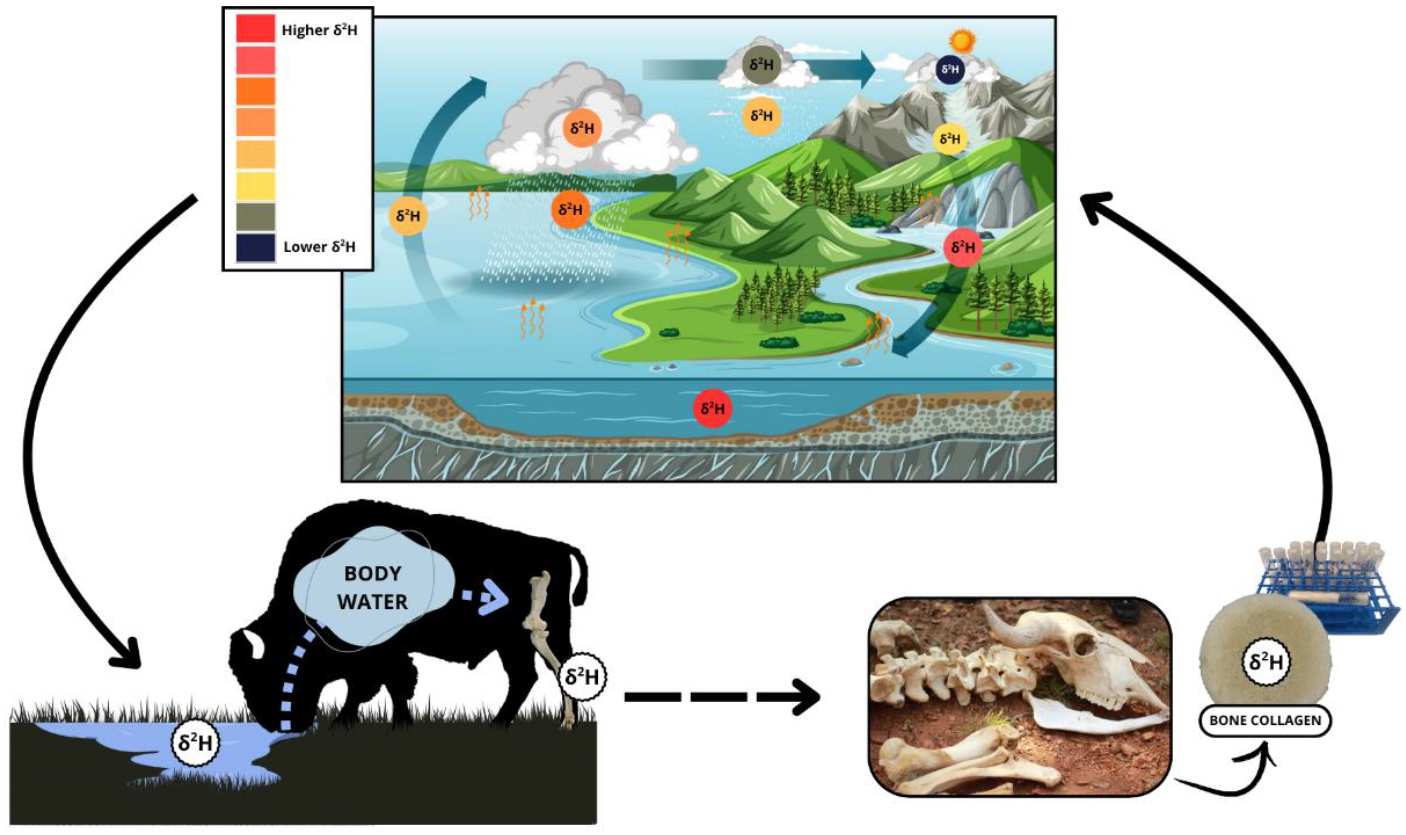
General outline of δ^2^H analysis approach, from the description of how the hydrological cycle generates a characteristic isotopic variation in δ^2^H, to how this value is subsequently assimilated into different animal tissues, and how the analysis of bone collagen samples can be used to interpret the climatic isotopic variations.

This study aims to investigate whether δ^2^H can serve as a proxy of local climatic evolution by leveraging the unique opportunities provided by archaeological deposits, where human groups accumulated faunal remains from the surrounding environment. To our knowledge, this is one of the first analyses of local δ^2^H evolution in bone collagen from Palaeolithic faunal remains over time. Our results demonstrate that hydrogen isotopes can be used to reconstruct general site-specific palaeoclimatic conditions and refine interpretations from previous palaeoenvironmental studies. Moreover, we identify species-specific δ^2^H differences, highlighting the need for further investigation into the physiological and ecological mechanisms influencing hydrogen isotope incorporation in animal tissues.

## 2. ARCHAEOLOGICAL SITE

Located at the confluence of Atlantic and continental climate influences, we have selected the Isturitz Cave (Western Pyrenees, France; *Figure 2A*) as a case study due to its well-documented archaeological record, which makes it particularly suitable for exploring climate variability over a controlled period of around 3000 years at a regional scale. Two galleries within the Isturitz karstic cave system have been excavated since 1912 (De Saint-Périer, 1935; Normand, 2005; Passemard, 1913), revealing a stratigraphic sequence spanning from the Mousterian to the Bronze Age (Normand et al., 2007).

**Figure 2.**
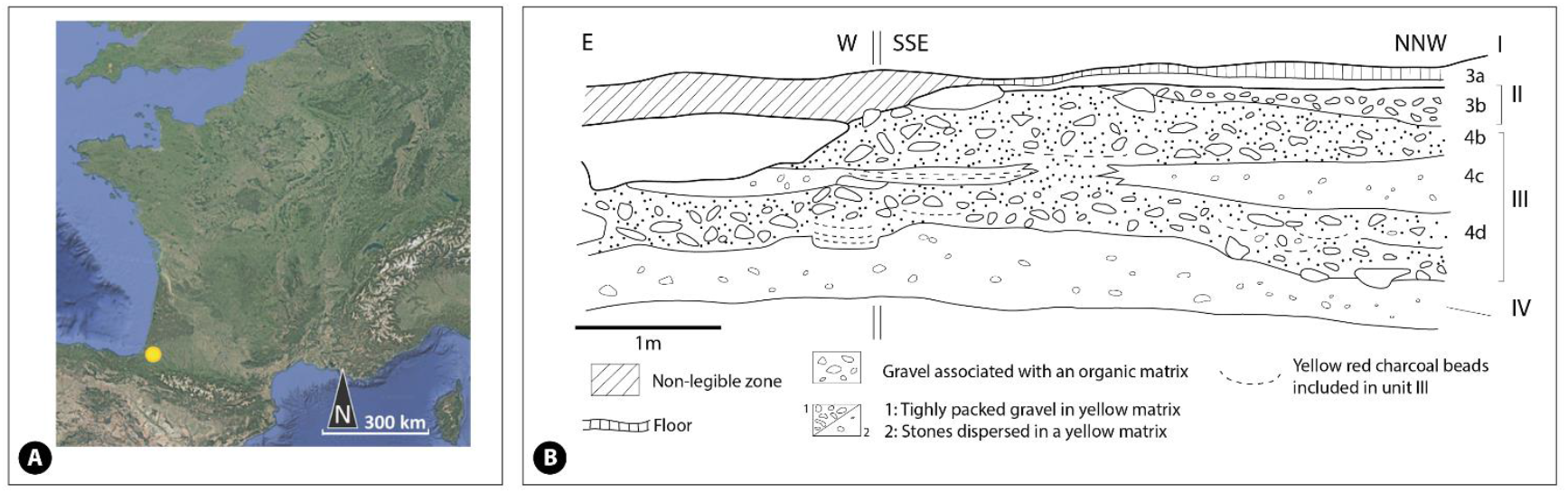
**A)** Location of the Isturitz cave in the valley of the river l’Arberoue in France (Western Pyrenees). Modified from Berlioz *et al*. (2025). **B)** Synthetic stratigraphy of the excavated zone, redraw from Normand *et al*. (2007).

We will focus on the Aurignacian occupation phases within lithostratigraphic unit III, which has revealed well-preserved archaeological materials (Soulier, 2013; Tarriño and Normand, 2002; White and Normand, 2015; *Figure 2B*). These Aurignacian levels are subdivided into three archaeostratigraphic phases—Proto-Aurignacian, Intermediate Aurignacian, and Early Aurignacian—originally dated by Barshay-Szmidt et al. (2018) and recently revised by Berlioz et al. (2025). The updated chronological framework places the Proto-Aurignacian between 42,470 and 41,460 cal BP, the Intermediate Aurignacian between 42,090 and 40,790 cal BP and the Early Aurignacian between 41,450 and 39,480 cal BP (hereinafter also referred to as Proto, Intermediate and Early). This means a temporal sequence between 42,470 and 39,480 cal BP.

## 3. MATERIALS AND METHODS

The carbon, nitrogen and sulphur isotopic composition of the samples used in this study were previously published by Berlioz et al. (2025). In order to complement the existing isotopic data, we conducted additional analysis targeting hydrogen isotopes once the following quality indicators were set in order to ensure collagen integrity: ≥ 1 % collagen (Ambrose, 1990; van Klinken, 1999), 2.9 – 3.6 C:N (Ambrose, 1990; M. DeNiro, 1985; Guiry and Szpak, 2021; van Klinken, 1999), 600 ± 300 C:S and 200 ± 100 N:S (Nehlich and Richards, 2009). We re-analysed a total of 50 bones of auroch/bison (*Bos primigenius/Bison priscus*, n=20; from now on named as bovine), horse (*Equus ferus*, n=20), and reindeer (*Rangifer tarandus*, n=10); eleven additional bones, including three red deer (*Cervus elaphus)*, seven fox (*Vulpes vulpes*) and one bear (*Ursus* sp.), were also analysed but excluded from further explorations because either species was present in only one cultural phase or in order to avoid dietary accumulation effects from carnivory (Birchall et al., 2005; Pietsch et al., 2011).

Bone samples were taken at the *Centre Départemental d’Archéologie of Hasparren* (France), then collagen extraction was undertaken at the EvoAdapta clean lab at the University of Cantabria (Santander, Spain) following the methodology detailed in Brown *et al*. (1988) and Richard and Hedges (1999). All isotopic analyses were performed by the Iso-Analytical Laboratory (Crewe, UK) using a Europa Scientific 20-20 isotope ratio mass spectrometer and the procedure detailed in Jones et al. (2019) was followed to measure δ^13^C, δ^15^N, and δ^34^S, as stated in Berlioz et al. (2025). Subsequently, the remaining collagen was prepared for δ^2^H analyses directly at Iso-Analytical facilities. As a precautionary measure, samples containing globules were freeze-dried before analysis to eliminate any possible water content. Samples were then weighed (1.0 ± 0.1 mg) into open silver capsules, alongside non-exchangeable hydrogen standards, and comparatively equilibrated with moisture in the laboratory air for 14 days prior to analysis. Once batch analysis had begun, the capsules were sealed and loaded into an auto-sampler, then dropped into a furnace at 1080 °C and thermally decomposed to H_2_ and CO over glassy carbon. Any traces of water produced were removed by magnesium perchlorate. H_2_ was resolved by a packed column gas chromatograph held at 35 °C. The resultant chromatographic peak entered the ion source of the IRMS, where it was ionised and accelerated. Gas species of different masses were separated in a magnetic field and then simultaneously measured on a Faraday cup universal collector array. For H_2_, masses 2 and 3 were monitored. Duplicate measurements were taken every five samples. Hydrogen isotopes quality measurements were checked using in-house standards IA-R002 (mineral oil: δ^2^H_V-SMOW_= -111.2 ‰) and IA-R072 (mineral oil: δ^2^H_V-SMOW_ = -148.61 ‰) that were calibrated against the International Atomic Energy Agency (IAEA) standard IAEA-CH-7 (polyethylene foil: δ^2^H_V-SMOW_ = -100.2 ‰). Deuterium non-exchangeable hydrogen measurements were comparatively equilibrated by a 3-point linear calibration and control checked by inter-laboratory United States Geological Survey (USGS) comparison standards: USGS42 (human hair: non-exchangeable δ^2^H_V-SMOW_ = -72.9 ‰), USGS43 (human hair: non-exchangeable δ^2^H_V-SMOW_ = -44.4 ‰), USGS CBS (caribou hoof: non-exchangeable δ^2^H_V-SMOW_ = -157.0 ‰) and USGS KHS (kudu horn: non-exchangeable δ^2^H_V-SMOW_ = -35.3 ‰). USGS43 was measured as a quality control check sample to verify the use of the 3-point linear calibration. Average reproducibility of samples measured in duplicate was 1.30‰ for δ^2^H and 1.51‰ for non-exchangeable δ^2^H. The δ^2^H, δ^13^C, δ^15^N, and δ^34^S values are reported relative to V-SMOW, V-PDB, AIR, and V-CDT standards.

Data was prepared and statistically analysed using R (R Core Team, 2024), RStudio v. 2024.12.1.563 (Posit team, 2025) and the packages “car” (Fox and Weisberg, 2019), “corrplot” (Wei and Simko, 2021), “dplyr” (Wickham et al., 2023), “FSA” (Ogle et al., 2023), “Hmisc” (Harrell Jr, 2025), “rstatix” (Kassambara, 2023), “tidyverse” (Wickham et al., 2019). For data groups with at least three stable isotopic values, the Kruskal-Wallis test was performed, followed by Dunn’s post-hoc analyses with Bonferroni adjustment when statistically significant differences were detected. All raw data and the R code file of the analysis are provided as Supplementary Information (S1 and S2 respectively).

## 4. RESULTS

### 4.1 Collagen preservation and sample selection

As noted above, of the 61 samples initially analysed, 50 exceeded all of our quality and methodological criteria. Therefore, all the samples that were further analysed had the expected terrestrial mammal values for preserved collagen, likely reflecting in vivo isotopic composition (Guiry and Szpak, 2021; Nehlich and Richards, 2009; van Klinken, 1999). The percentage of collagen for them ranged from 2.15 to 10.04%, with an average of 5.74%; for the C:N the range was 3.11 to 3.22 with an average of 3.15; for the C:S ratio, the minimum value was 322.99 and the maximum 729.22, while the average was 536.98; finally, for the N:S values, the minimum was 100.51 and the maximum 230.89, with an average of 170.59. Reindeer samples were discarded for interspecific statistical analysis because they did not meet the minimum number of individuals threshold (n ≥3).

### 4.2 δ^2^H, δ^13^C, δ^15^N, and δ^34^S isotope data

Main isotopic results are shown in *Table 1*. Considering all herbivores species and cultural periods together, we found a mean value of –20.59 ± 0.77‰ for δ^13^C, of 7.18 ± 1.99‰ for δ^15^N, of 12.61 ± 1.4‰ for δ^34^S and of –51.24 ± 11.51‰ for δ^2^H_non-exchangeable_. As expected, when evaluating the relationship between the different isotopes, we found a strong positive correlation between δ^13^C and δ^15^N (Pearson’s r= 0.64, P=<0.001, *Figure 3*). There was also a strong negative correlation between δ^2^H_non-exchangeable_ and δ^13^C (Pearson’s r =-0.37, P=0.007) and between δ^2^H_non-exchangeable_ and δ^15^N (Pearson’s r= -0.54, P=<0.001).

**Table 1.**
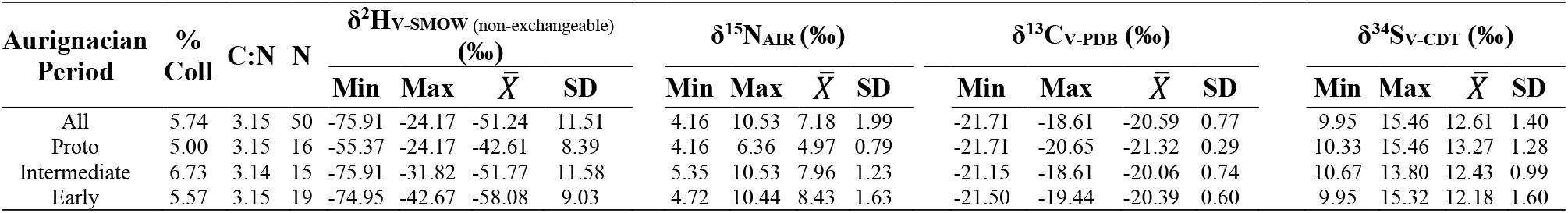
Results of isotope data (δ^13^C, δ^15^N, δ^34^S and δ^2^Hnon-exchangeable) by periods from bone collagen of Isturitz. Abbreviations: % Coll = mean percentage of extracted collagen; C:N = mean ratio of carbon (δ^13^C) and nitrogen (δ^15^N); N = sample size; Min. = minimum value; Max. = maximal value; 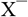 = mean value; SD = Standard Deviation.

| Aurignacian Period | % Coll | C:N | N | $\delta^2H_{\text{V-SMOW (non-exchangeable)}}$ (‰) | | | | $\delta^{15}N_{\text{AIR}}$ (‰) | | | | $\delta^{13}C_{\text{V-PDB}}$ (‰) | | | | $\delta^{34}S_{\text{V-CDT}}$ (‰) | | | |
| --- | --- | --- | --- | --- | --- | --- | --- | --- | --- | --- | --- | --- | --- | --- | --- | --- | --- | --- | --- |
| | | | | Min | Max | $\bar{X}$ | SD | Min | Max | $\bar{X}$ | SD | Min | Max | $\bar{X}$ | SD | Min | Max | $\bar{X}$ | SD |
| All | 5.74 | 3.15 | 50 | -75.91 | -24.17 | -51.24 | 11.51 | 4.16 | 10.53 | 7.18 | 1.99 | -21.71 | -18.61 | -20.59 | 0.77 | 9.95 | 15.46 | 12.61 | 1.40 |
| Proto | 5.00 | 3.15 | 16 | -55.37 | -24.17 | -42.61 | 8.39 | 4.16 | 6.36 | 4.97 | 0.79 | -21.71 | -20.65 | -21.32 | 0.29 | 10.33 | 15.46 | 13.27 | 1.28 |
| Intermediate | 6.73 | 3.14 | 15 | -75.91 | -31.82 | -51.77 | 11.58 | 5.35 | 10.53 | 7.96 | 1.23 | -21.15 | -18.61 | -20.06 | 0.74 | 10.67 | 13.80 | 12.43 | 0.99 |
| Early | 5.57 | 3.15 | 19 | -74.95 | -42.67 | -58.08 | 9.03 | 4.72 | 10.44 | 8.43 | 1.63 | -21.50 | -19.44 | -20.39 | 0.60 | 9.95 | 15.32 | 12.18 | 1.60 |

**Figure 3.**
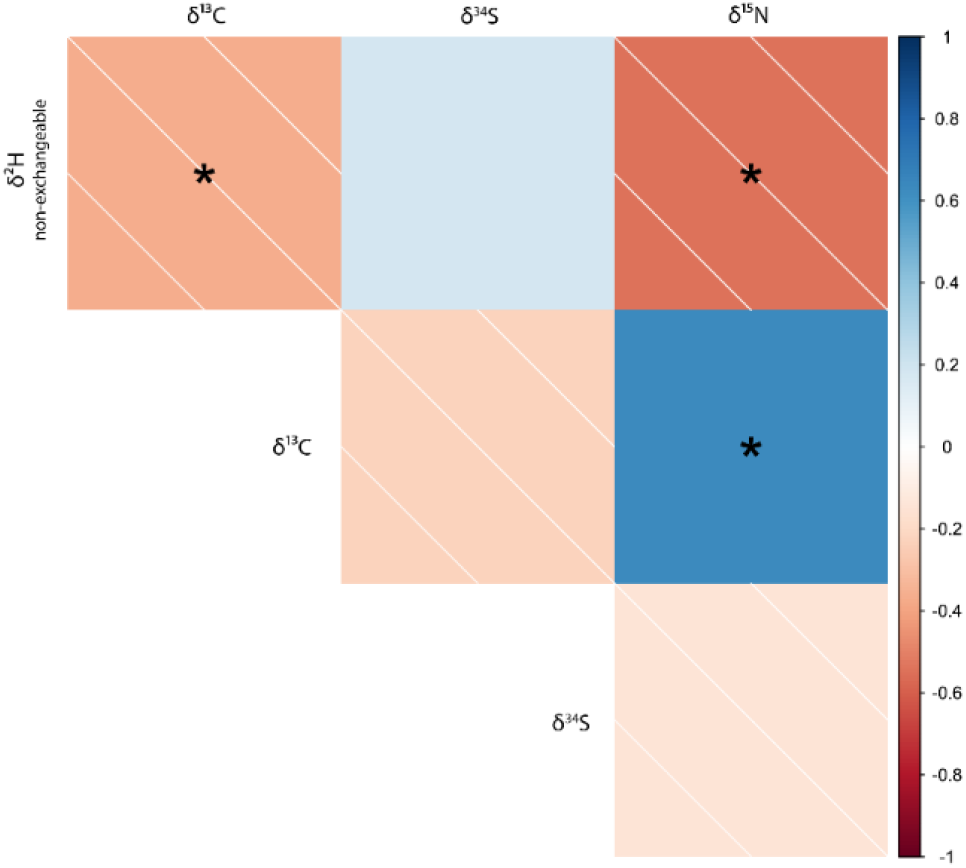
Correlation matrix showing Pearson’s r between the analysed stable isotope ratios (‰). Significant correlations (p < 0.05) are marked with asterisks (*).

Considering the data from all species together, it seems that the δ^13^C evolution through the Aurignacian started with an increase from the Proto to the Intermediate period (δ^13^C_Proto_=-21.32 ± 0.29‰, and δ^13^C_Intermediate_=-20.06 ± 0.74‰), followed by a small decrease in the Early period (δ^13^C_Early_=-20.39 ± 0.60‰). However, even though the Kruskal-Wallis test revealed significant differences for this isotope (ꭓ^2^=28.58, P=<0.001), Dunn’s post-hoc test showed that only the transition from the Proto-Aurignacian to the Intermediate Aurignacian presents significant differences (z=4.02, P=<0.001; see *Table 2* and *Figure 4*).

**Table 2.** Kruskal-Wallis test and Dunn’s post-hoc on the variability of all analysed isotopes for each archaeostratigraphic phase. The z-score correspond to the difference in the mean variability between the compared groups. When Kruskal-Wallis was not significant (p.value >0.05), Dunn’s post-hoc were not performed. Bold type indicates significant differences in mean variability between periods.

| Stable isotope | Kruskal-Wallis |  |  | Dunn's post-hoc |  |  |
| --- | --- | --- | --- | --- | --- | --- |
| | $\chi^2$ | DF | p-value | Aurignacian Period | z-score | p-value |
| $\delta^2\text{H}_{\text{(non-exchangeable)}} (\text{‰})$ | 16.826 | 2 | **<0.001** | Proto – Intermediate | -2.093 | 0.109 |
|  |  |  |  | Proto – Early | -4.102 | **<0.001** |
|  |  |  |  | Intermediate – Early | -1.852 | 0.192 |
| $\delta^{15}\text{N} (\text{‰})$ | 26.454 | 2 | **<0.001** | Proto – Intermediate | 4.815 | **<0.001** |
|  |  |  |  | Proto – Early | 4.021 | **<0.001** |
|  |  |  |  | Intermediate – Early | -1.06 | 0.868 |
| $\delta^{13}\text{C} (\text{‰})$ | 28.575 | 2 | **<0.001** | Proto – Intermediate | 4.016 | **<0.001** |
|  |  |  |  | Proto – Early | 5.088 | **<0.001** |
|  |  |  |  | Intermediate – Early | 0.819 | 1.000 |
| $\delta^{34}\text{S} (\text{‰})$ | 5.715 | 2 | 0.058 | - | - | - |

**Figure 4.**
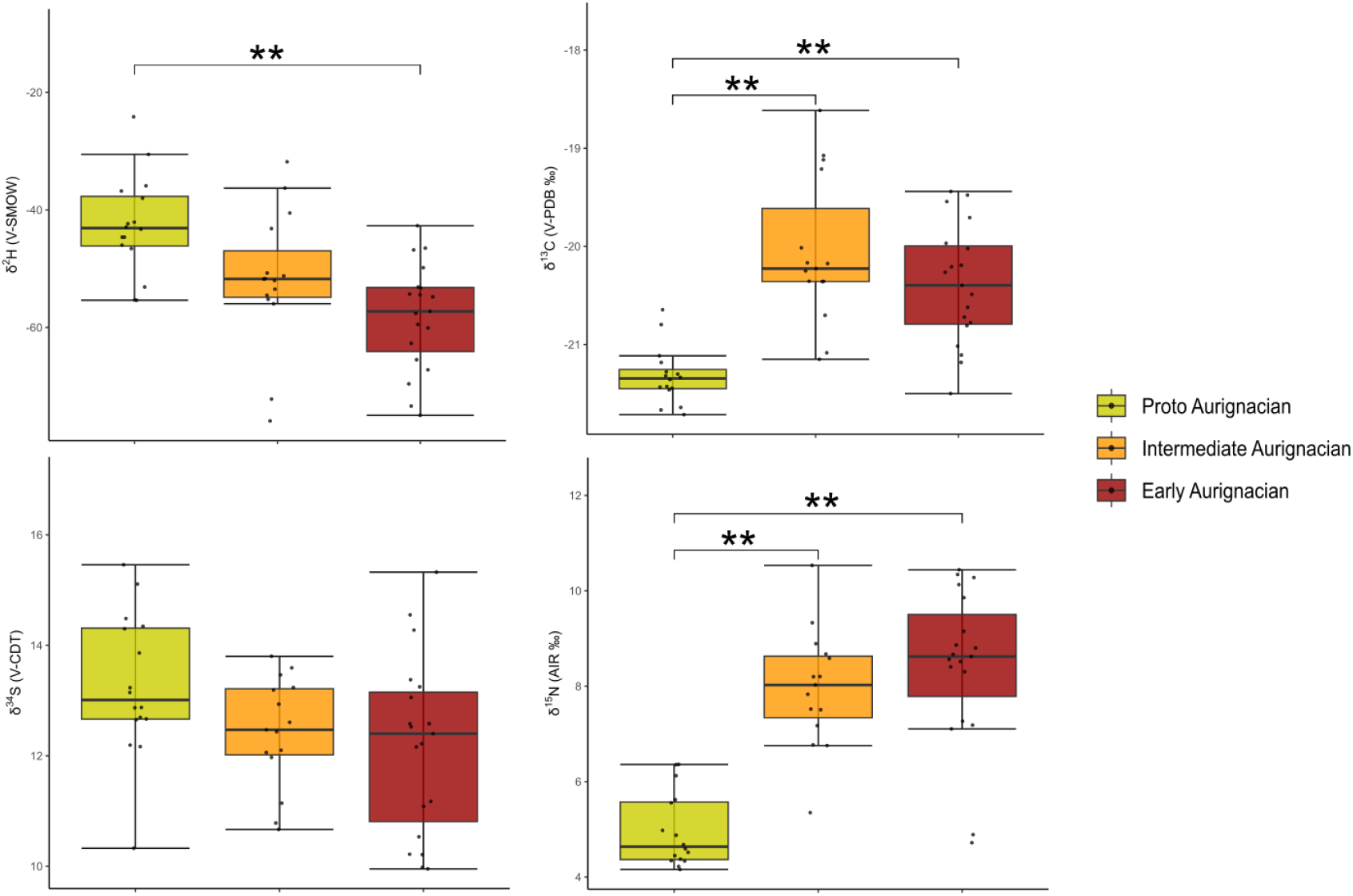
δ^13^C, δ^2^H, δ^34^S and δ^15^N values from the bone collagen of Isturitz (mean ± SD). Significant differences (p < 0.05) are marked with asterisks (** refers to p-values < 0.001).

Conversely, δ^15^N seemed to increase over time (δ^15^N_Proto_=4.97 ± 0.78‰, δ^15^N_Intermediate_=7.96 ± 1.23‰ and δ^15^N_Ancien_=8.43 ± 1.63‰), but again, the only successive period change that was significant was that from the Proto to the Intermediate Aurignacian (Kruskal-Wallis test: (ꭓ^2^=26.45, P=<0.001; Dunn’s test: z=4.02, P=<0.001; Table 2). Hydrogen showed the opposite behaviour, apparently decreasing with time (δ^2^H_non-exchangeable∼Proto_=-42.61 ± 8.39‰, δ^2^H_non-exchangeable∼Intermediate_=-51.77 ± 11.58‰ and δ^2^H_non-exchangeable∼Ancien_=58.08 ± 9.03‰); in this case, the only significant difference takes place between the Proto-Aurignacian and the Early Aurignacian (Kruskal-Wallis test: ꭓ^2^=16.83, P=<0.001; Dunn’s test: z=-4.10, P=<0.001; Table 2, *Figure 4*). For δ^34^S, we found a similar visual decreasing behaviour (δ^34^S_Proto_=13.27 ± 1.28‰, δ^34^S_Intermediate_=12.43 ± 0.99‰ and δ^34^S_Ancien_=12.18 ± 1.60‰), but there was no significant difference between the archaeological stratigraphic units (ꭓ^2^=5.71, P=0.06).

### 4.3 Interspecific isotope data

The results, categorised by species, are presented in *Table 3*. The three species *E. ferus, R. tarandus* and *B. primigenius/Bi. priscus* presented similar mean values for δ^13^C (*R. tarandus*=–19.66 ± 0.87‰; *Bovines*=-20.54 ± 0.54‰; *E. ferus*=-21.05 ± 0.47‰), δ^15^N (*R. tarandus*=7.68 ± 1.46‰; *Bovines*=7.45 ± 1.85‰; *E. ferus*=6.76 ± 2.31‰) and δ^34^S (*R. tarandus*=11.72 ± 1.15‰; *Bovines*=13.08 ± 1.17‰; *E. ferus*=12.47 ± 1.52‰). Non-exchangeable hydrogen also showed similar values between species (*R. tarandus*=-50.86 ± 6.88‰; *Bovines*=-50.58 ± 14‰; *E. ferus*=-52.09 ± 13.95‰), although it was noticeable that the variability between samples is higher than for the rest of the isotopes analysed.

**Table 3.**
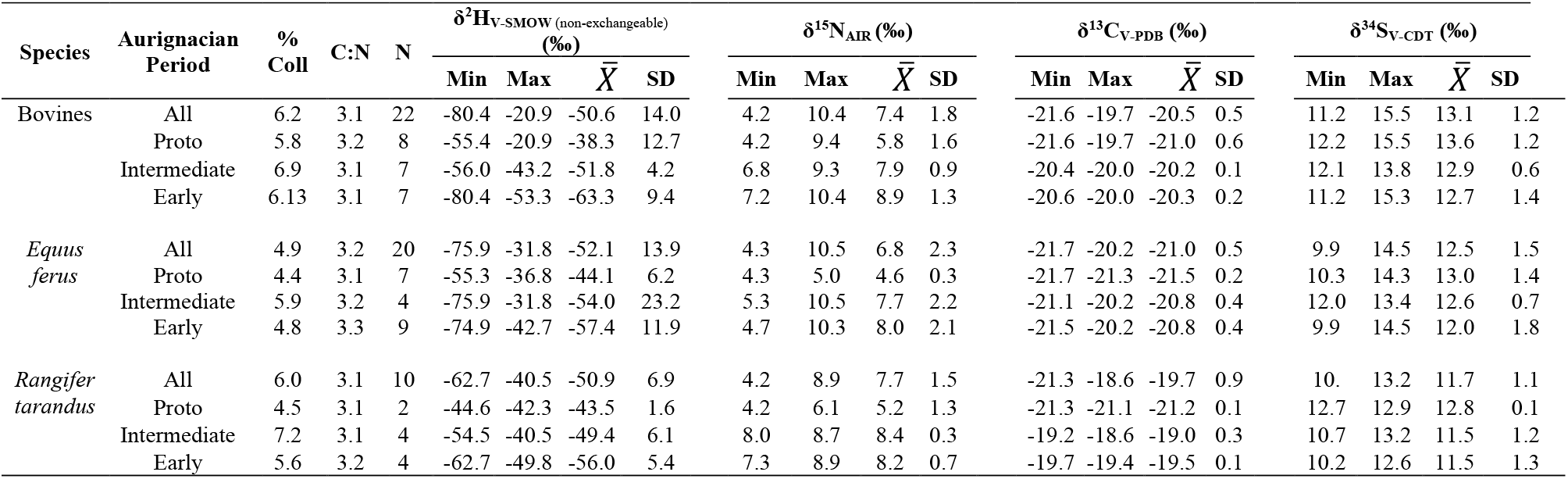
Results of isotope data (δ^13^C, δ^15^N, δ^34^S and δ^2^H_non-exchangeable_) by species from bone collagen of Isturitz. Abbreviations: % Coll = mean percentage of extracted collagen; C:N = mean ratio of carbon (δ^13^C) and nitrogen (δ^15^N); N = sample size; Min. = minimum value; Max. = maximal value; 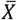 = mean value; SD = Standard Deviation.

The results of the statistical analyses indicate that none of the species studied showed significant differences in any of the archaeological periods in terms of δ^34^S (ꭓ^2^_*R*.*tarandus*_=0.33, P_*R*.*tarandus*_=0.56; ꭓ^2^_Bovines_=1.73, P_Bovines_=0.42; ꭓ^2^_*E*.*ferus*_=1.81, P_*E*.*ferus*_=0.56; see *Table 4*). As previously reported by Berlioz et al. (2025), there were significant differences between the Proto-Aurignacian and both the Intermediate and the Early Aurignacian for δ^15^N in *B. primigenius/Bi. priscus* (z_Early-Proto_=3.65, P=<0.001; z_Intermediate-Proto_=2.62, P=0.03) and *E. ferus* (z_Early-Proto_=3.17, P= < 0.01; z_Intermediate-Proto_=2.45, P=0.04), where values appeared to increase over time (*Figure 5*). Same pattern was also described for δ^13^C levels, not only in bovines (z_Early-Proto_=2.84, P=0.01; z_Intermediate-Proto_=3.34, P=<0.01) and horses (z_Early-Proto_=2.85, P=0.01; z_Intermediate-Proto_=2.64, P=0.02); but also, from the Intermediate to the Early Aurignacian in reindeer, the only transition that could be studied for this species and therefore was only analyzed by Kruskal-Wallis test (ꭓ^2^=5.33, P=0.02).

**Table 4.**
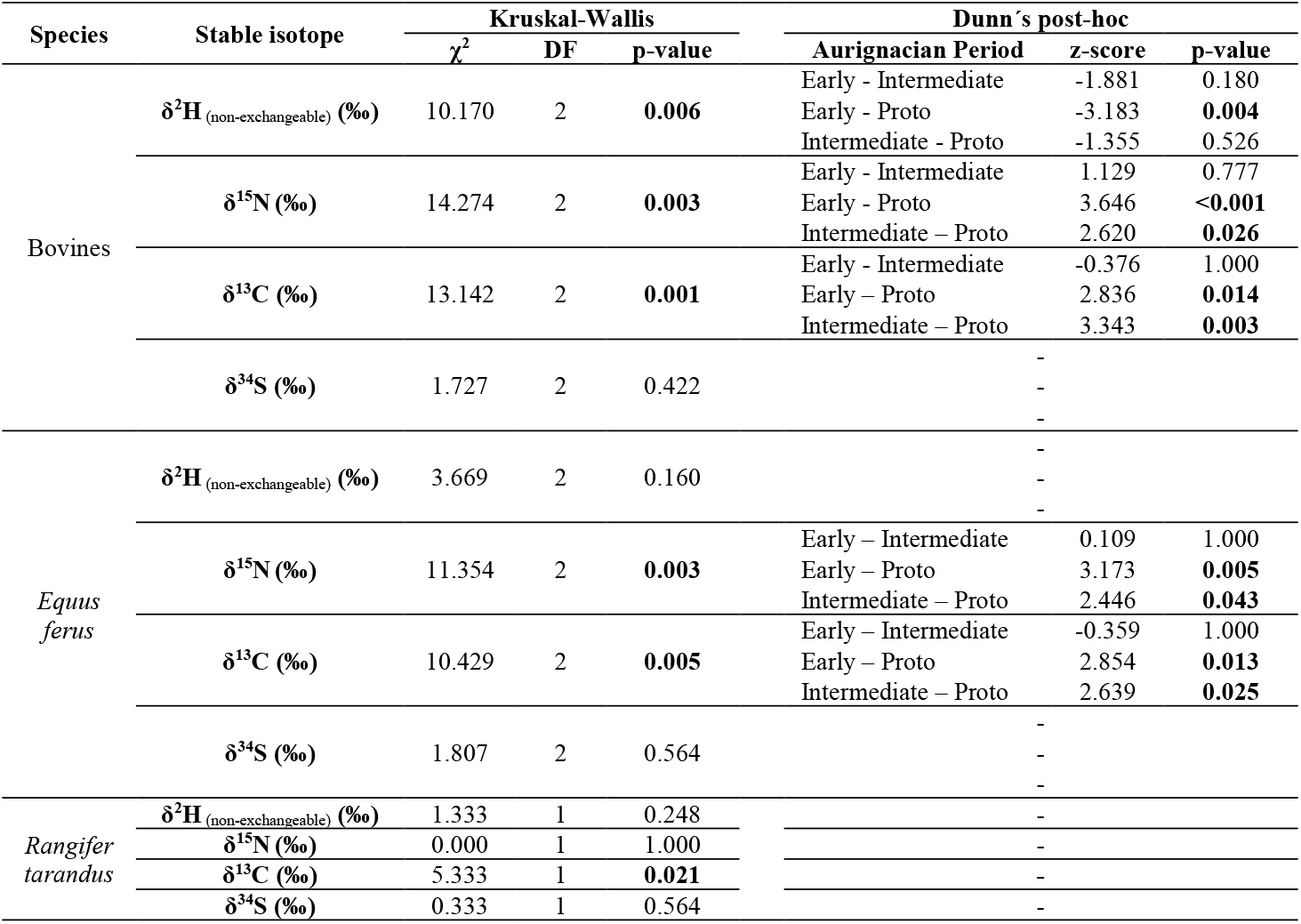
Kruskal-Wallis test and Dunn’s post-hoc on the variability of all analysed isotopes for each species and archaeostratigraphic phase. The z-score correspond to the difference in the mean variability between the compared groups. When Kruskal-Wallis is not significant (p.value >0.05), Dunn’s post-hoc are not reported. Bold type indicates significant differences in mean variability between periods; except for *Rangifer tarandus*, for which no such analysis was carried out.

**Figure 5.**
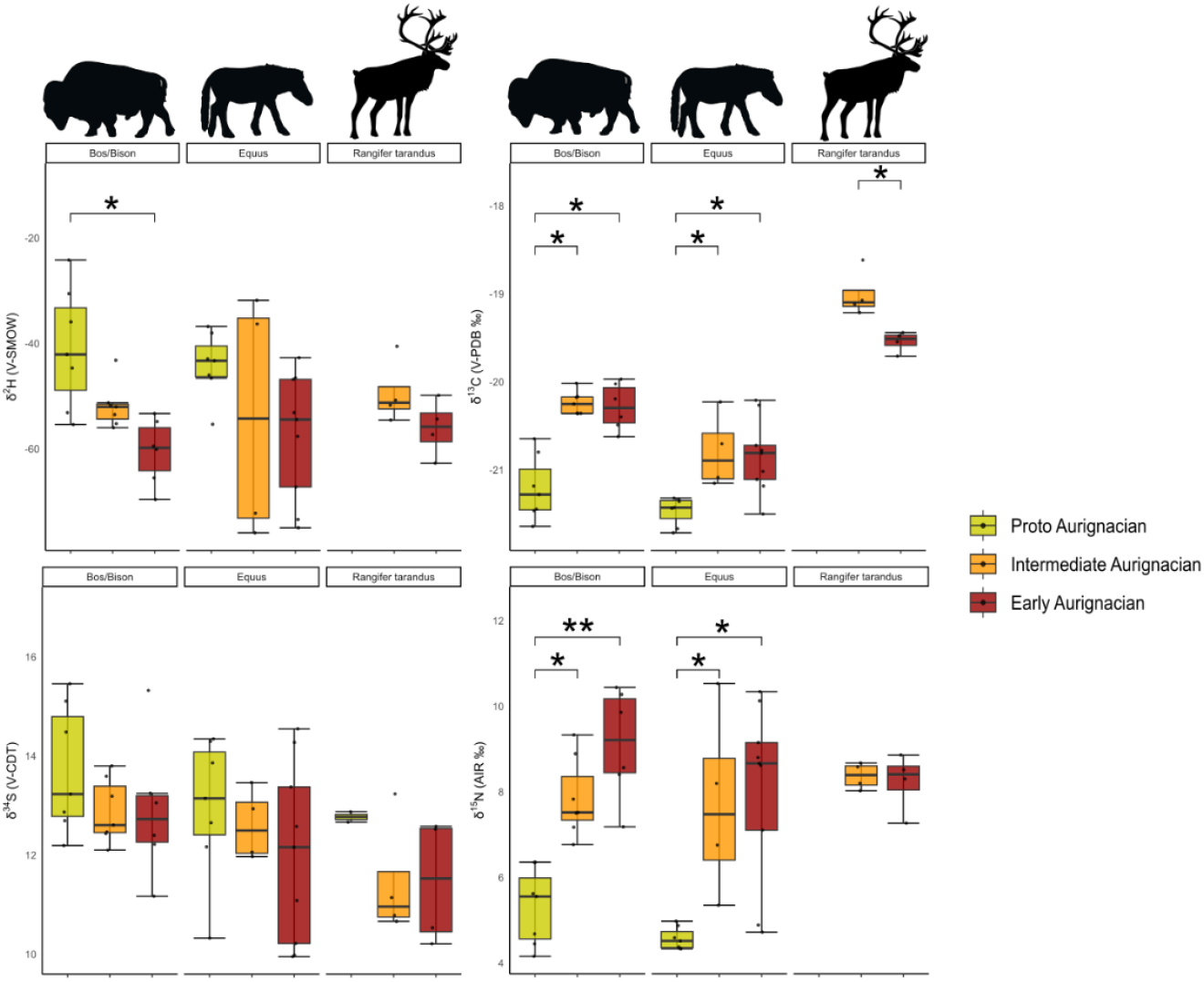
δ^13^C, δ^2^H, δ^34^S and δ^15^N values from the bone collagen of Isturitz (mean ± SD) by species. Significant differences (p < 0.05) are marked with asterisks (* refers to p-values <0.05, while ** indicates values < 0.001).

As for δ^2^H, we found fewer statistical differences. Only the values between the Proto-Aurignacian and the Early Aurignacian of bovids differed significantly, decreasing from the oldest to the most modern period (z_Early-Proto_=-3.18, P=0.004).

## 5. DISCUSSION

Reconstructing palaeoclimatic conditions remains a methodological challenge, often constrained by the nature of available materials and proxies. In this study, we experimentally tested the use of δ^2^H in bone collagen dated between 42,470 and 39,480 cal BP as a novel isotopic approach to obtain past local climatic information. Our results reveal significant variation in δ^2^H values across the Aurignacian sequence at Isturitz, broadly consistent with expected changes in precipitation. However, species-specific variation suggests physiological and ecological factors also influence the incorporation of δ^2^H into collagen.

The isotopic composition of biological tissues is to some extent a reflection of feeding behaviour, and hydrogen would be no exception. While the degree to which δ^2^H values originate from diet or drinking water is still debated, the most pronounced dietary effects are typically observed across trophic levels, where it is potentially valuable for differentiating between more herbivore or carnivore diets, even among omnivores (Birchall et al., 2005; Clauzel et al., 2022; Gröcke et al., 2017; Hobson et al., 1999; Newsome et al., 2020; Reynard et al., 2025). As our analysed dataset only considers herbivores, this possible trophic bias is minimised. Thus, the observed trend— from higher δ2H values in the Proto-Aurignacian to lower values in the Early Aurignacian — is more likely a reflection of changes in drinking water sources. Hydrogen isotopes values in the water cycle depend on local factors such as distance to the sea, altitude and amount of precipitation, among others (Dansgaard, 1964; Dee et al., 2023; Kern et al., 2020; Pfahl and Sodemann, 2014). Hence, the change we observed in our results could be driven not only by climatic factors, but also by differences in the location from which the analysed species obtained their water. We analysed the δ^34^S in order to address this duality, considered as a geographical proxy for its mainly dependence on geological composition and sea spray (Drucker et al., 2011; Guiry and Szpak, 2021). The absence of significant differences in sulphur over time suggests that the same landscape was exploited throughout the entire sequence. This supports the interpretation that the observed hydrogen isotopes variations are a reflection of evolving local climatic conditions, such as rainfall regimes. Indeed, similar correlations between precipitation and collagen δ^2^H have been reported in modern and archaeological herbivores (Cormie et al., 1994; Leyden et al., 2006; Reynard et al., 2020; Topalov et al., 2013).

Further support for a progressive aridification is provided by Berlioz and collaborators (2025), who combined δ^18^O and δ^13^C from tooth enamel and δ^13^C and δ^15^N from bone collagen to obtain a climatic reconstruction of Isturitz. Their results suggest a decrease in both humidity and temperature over time from the Proto-to the Early Aurignacian. However, δ^13^C and δ^15^N reflect environmental proxies, not direct climatic variables, and are more sensitive to changes in vegetation and dietary behaviour. In fact, even though we have found the expected correlations between carbon and nitrogen isotopes and hydrogen, the absorption of these isotopes varies across different periods. Besides the change we observed from the beginning to the end of the Early Aurignacian for the three isotopes (δ^2^H, δ^13^C and δ^15^N), we also observed a significant shift in δ^13^C and δ^15^N from the Proto-Aurignacian to the Intermediate stage. Berlioz et al. (2025) interpreted this shift as an abrupt drop in precipitation and temperature, followed by slight improvements in both variables during the Early Aurignacian. The absence of a parallel shift in δ^2^H, however, suggests that precipitation patterns changed more gradually, and thus has a delayed or buffer effect on drinking water hydrogen values.

Environmental changes in the plant community could partly explain this discrepancy. The levels of δ^13^C and δ^15^N in plants vary due to multiple factors, including climatic conditions and the morphology and physiology of each species. Our carbon isotopic values reflect the dominance of C_3_ plants in open landscapes across the sequence, consistent with Berlioz and collaborators (2025). Therefore, a change in the canopy on C3 plants would not be the primary driver of this isotopic shift. Another possibility could be the species of plants consumed by the analysed herbivores. Palynological analysis of Isturitz samples showed a change in the representation of plant biodiversity from *Pinus, Alnus, Asteraceae* and *Poaceae* during the Intermediate Aurignacian, to an increase in the variety of represented taxa during the Early Aurignacian, including species such as *Betula, Corylus* and *Quercus*, as well as the already known *Pinus, Alnus, Asteraceae* and *Poaceae* (Normand et al., 2007). No pollen analysis has been conducted specifically on the Proto-Aurignacian of Isturitz, nor during the excavations of C. Normand or A. Leroi-Gourhan (Leroi-Gourhan, 1959; Normand et al., 2007). However, the results of marine core analysis in the Bay of Biscay pointed to the general abundance of Atlantic forests and semidesert steppes in the region during this period based on the presence of pollen from *Betula*, deciduous *Quercus, Pinus, Alnus, Corylus, Carpinus, Calluna, Ericaceae, Artemisia, Amaranthaceae, Cyperaceae*, and *Poaceae* (Fourcade et al., 2022; Sánchez Goñi et al., 2008). This pattern suggests that woody species temporarily retreated under drier conditions and expanded again with either a slight climatic improvement, even in the absence of major increases in precipitation, or because of the stabilisation of better-adapted plant communities (Zhang et al., 2023). Such vegetation dynamics would be consistent with a moderate but continuous aridification that may not be fully captured in collagen δ^2^H values.

Moreover, the significant increase in both δ^13^C and δ^15^N values between the Proto and Intermediate Aurignacian may reflect not only a shift in biodiversity but also specific adaptations in plant carbon and nitrogen fixation under drought stress. During the early stages of moderate but sustained aridification, C_3_ plants can enhance carbon assimilation and water-use efficiency through stomatal regulation and biochemical adjustments (Robertson et al., 2021; Wittmer et al., 2008). These responses can lead to slightly enriched δ^13^C values in plant tissues, which would be transmitted into the collagen of herbivores that consume them. A similar enrichment in δ^15^N may also occur, though this signal is more often linked to shifts in nitrogen sources and uptake strategies under arid conditions than to direct metabolic enhancement (Amundson et al., 2003; Hartman and Danin, 2010). As drought intensifies, species with fibrous root systems and rapid stomatal regulation are likely to gain dominance, outcompeting less drought-tolerant taxa. Such ecological turnover could stabilize isotopic values despite ongoing environmental stress, thereby explaining the absence of significant isotopic changes between the Intermediate and the Early Aurignacian. Furthermore, such physiological responses are often species-specific. For example, *Pinus* species often show elevated δ^13^C values under drought conditions, likely due to more conservative or controlled stomatal behaviour and a lower discrimination towards this isotope (Diefendorf et al., 2010; Martínez-Sancho et al., 2018). This physiological adaptation locates *Pinus* between early-successional species during the initial stages of ecosystem recovery after periods of disturbance. Therefore, its persistence not only throughout the entire Aurignacian period, but especially after the first stages of aridification during the Intermediate Aurignacian, is not surprising (Beloiu Schwenke et al., 2024). The same applies to the genus *Alnus*, whose species are known to be symbiotically related to the actinobacterium *Frankia*, being typical nitrogen-fixing non-leguminous trees. In general, both soil and plant δ^15^N values increase systematically along aridification, but this would be even more noticeable across time in species like this, which under stress conditions change the main assimilation of nitrogen from their actinorhizal symbiosis to nitrogen from soil (Hartman and Danin, 2010; Kohls et al., 2003). Taken together such physiological and ecological dynamics would be expected to influence the isotopic composition of herbivore bone collagen. Individuals with narrow dietary breadth may show stronger reflection of these plant-level adaptations, especially where Pinus and Alnus, were prevalent during the Intermediate Aurignacian, whereas animals with access to a more diverse plant community would exhibit more variable isotopic signatures. Beyond these species-specific examples, the broader implication is that plant physiology and animal dietary behaviour must be considered when interpreting isotopic data. The absence of a corresponding shift in δ^2^H supports the scenario of gradual aridification, as hydrogen isotopes are less influenced by trophic or metabolic factors. This multi-isotopic approach highlights both, the complexity of ecosystems and the challenges of making direct inferences about climate through values that are more closely related to environmental factors.

Likewise, when interpreting isotopic data from bone collagen, it is essential to account for interspecific differences in feeding strategies and water use, as these directly influence the isotopic signals preserved in tissues. Based on carbon and nitrogen isotope results, Berlioz and collaborators (2025) found that horse, bovids and reindeer generally exhibited grazing behaviour, consistent with dental microwear analyses. However, they also noted individual-level variability indicative of opportunistic or flexible feeding strategies, potentially associated with the use of woodland habitats. This variation is particularly evident in carbon isotopes. Reindeer, for example, tend to exhibit elevated δ^13^C values compared to other herbivores due to lichen consumption, which is also linked to seasonal grazing and aridification conditions at the intraspecific level (Takken Beijersbergen et al., 2021). Although our dataset for reindeer is limited, both in sample size and temporal representation, the fact that we not only observed such interspecific elevated values but also a significant change in the transition from the Intermediate to the Early Aurignacian only for this isotope, would again suggest and highlight how this isotope is more reflective of the environmental exploitation performed by these animals. By the same token, under cooling conditions, when preferred forage became scarce, Bisons can also incorporate lichens and mosses into their diet as fallback resources (Hecker et al., 2021; Julien et al., 2012; Jung, 2015). This strategy would also have a significant impact on carbon isotope values and may help explain the pronounced increase in δ13C observed in the species between the Proto- and Intermediate Aurignacian, a pattern not equally mirrored in equids. Besides diet alone, species-specific behaviours -such as feeding selectivity, plant part preference, seasonal mobility, and even sex-related differences-can influence both δ^13^C and δ^15^N values (Berini and Badgley, 2017; Britton et al., 2023; Metcalfe, 2021). These behavioural and physiological variables likely contribute to the significant shifts in carbon and nitrogen isotopes we observed between the Proto- and the Intermediate Aurignacian for bovids and horses. The consistency of the patterns across species suggests niche partitioning or specialization is unlikely to explain the observed isotopic variability.

Given this, it is also reasonable to consider that δ^2^H values could be influenced by species-specific behaviour in water acquisition and metabolism. Indeed, in our dataset, a significant shift in δ^2^H was recorded only for *Bos/Bison*. Among the species analysed, the absence of change in reindeer would be expected, as they possess highly efficient water conservation strategies -including moisture extraction from vegetation and snow, and reduce respiratory water loss-which enable them to survive and thrive in cold environments (Dryden, 2011; Nilssen et al., 1990). This ethological behaviour could mask the precipitation-derived hydrogen isotopes signal in tissues.

Horses and bovids, by contrast, are considered obligate drinkers and depend on regular access to free water. Yet only bovids exhibited a clear δ^2^H shift through the Aurignacian sequence. This discrepancy may be related to differences in physiology and water flux. Horses rely heavily on sweating for thermoregulation, which increases evaporative loss and may influence the fractionation of body water isotopes in these animals, thereby obscuring the climatic signal of δ^2^H in collagen (Arnold et al., 2006; Scheibe et al., 1998). Bovids, on the other hand, retain water more efficiently, and their larger body sizes results in higher body water volume and greater water turnover, making them more likely to reflect the isotopic signature of drinking water more accurately (Magozzi et al., 2019). Building on previous species-specific results regarding bone collagen hydrogen isotopes (Reynard et al., 2020), these findings suggest that bovids may serve as more reliable indicators of local hydrological conditions, while species such as horses and reindeer require more cautious interpretation.

Overall, our results reveal clear shifts in hydrogen values in bone collagen over 3,000-year time span, thereby indicating the climatic changes that modern humans faced during their occupation of the Aurignacian at Isturitz. This finding highlights the potential of δ^2^H in bone collagen as a proxy for reconstructing local palaeoclimatic conditions, particularly precipitation regimes, as well as attesting to its usefulness in Palaeolithic contexts when achieved on ungulates with evidence of human manipulation, such as the butchering process. While traditional isotopes such as δ^13^C and δ^15^N remain valuable indicators of environmental change, their interpretation is often complicated by dietary and ecological variables. In contrast, δ^2^H appears less affected by such factors in obligate drinkers, such as bovids, offering a more direct signal of hydrological change. The integration of δ^2^H with established isotopic systems provides a more nuanced understanding of past climate-environment-faunal interactions. Importantly, the faunal remains analysed here derive from archaeological human occupation levels, which introduces an inherent selection bias related to past hunting strategies and site-occupation patterns. While this limits the representativeness of local faunal communities, it also provides a valuable lens through which to explore how prehistoric humans interacted with and responded to changing environments. The integration of δ^2^H with established isotopic systems thus not only refines palaeoclimatic reconstructions but also offers new opportunities to investigate the ecological context of human behaviour. Future studies incorporating broader taxonomic samples and higher temporal resolution will be essential to validate δ^2^H as a robust palaeoclimatic proxy, to reveal its isoscapes and to further unravel the complex interplay between climate, ecosystems, and human adaptation.

## 6. CONCLUSIONS

This study provides one of the first assessments of δ^2^H in bone collagen from Palaeolithic faunal remains as a proxy for local climatic conditions. The significant variations in δ^2^H values observed across the Aurignacian sequence at Isturitz suggest a gradual decline in precipitation, broadly consistent with reconstructions based on other isotopic and palaeoenvironmental indicators. The absence of parallel shifts in δ^34^S supports animals stable inhabitation of the area used as hunting grounds by humans over time, strengthening the interpretation that δ^2^H changes reflect evolving local climate rather than geographic shifts in resource exploitation.

Species-specific responses highlight the influence of physiological and behavioural factors on isotope incorporation, particularly in relation to water intake and dietary preferences. Notably, δ^2^H shifts were recorded only in bovids, supporting their utility as reliable indicators of climate. By contrast, more complex isotopic responses in δ^13^C and δ^15^N appear driven by changes in plant physiology under drought stress and the dietary ecology of different herbivores.

As these faunal samples come from anthropogenic levels within a significant archaeological site, they inevitably reflect human hunting choices and seasonal occupation patterns. While this introduces potential biases in species representation, it also offers valuable insights into how Upper Palaeolithic human groups may have adapted to changing climatic and ecological conditions. Integrating δ^2^H with other isotopic and climatic proxies in archaeological contexts opens promising avenues for exploring both past climates and human-environment interactions in increasingly nuanced ways.

## AUTHOR CONTRIBUTIONS

A.B.M.-A. conceived the study, obtained funding to carry out the research, and together with L.A.-P. and M.F.-G. designed the research approach. L.A.-P. and A.C.-C. performed the majority of laboratory analyses. M.I.-H. performed the data processing and analysis. L.A.-P., M.F.-G. and M.C.S. contributed to the sampling process. C.N. and A.B.M.-A. provided the archaeological context of the Isturitz cave assemblage. M.I.-H. drafted the initial version of the manuscript, and all authors contributed to revisions and approved the final version of the text.

## DATA AVILABILITY

***Supplementary 1 (S1):*** Raw sample information for bone collagen samples and Hydrogen, Nitrogen, Carbon and Sulphur results with quality indicators.

***Supplementary 2 (S2):*** R code used for data processing and statistical analyses.

## AKNOWLEDGMENTS

Funding was mainly provided by the NEWINDS (Ref. PID2021-125818NB-I00) project from the Spanish Ministry of Science, Innovation and Universities to A.B.M-A, which also supports M.I-H through an FPI-PhD contract (Ref. PRE2022-105350); and by SUBSILIENCE (ID: ERC CoG Ref. 818299), which supported M.F.-G. through a postdoctoral contract during the data collection process, and who is currently supported by an APOSTD postdoctoral fellowship (CIAPOS/2022/081), funded by the Generalitat Valenciana and the European Social Fund. We gratefully acknowledge the sampling made by G. Terlato, hired within the SUBSILIENCE project; as well as Joëlle Darricau (owner of the archaeological site) and Olivier Ferullo (IE DRAC Nouvelle-Aquitaine) for making the prehistoric materials available to research.

